# Reconstruction of Gene Regulatory Networks by integrating biological model and a recommendation system

**DOI:** 10.1101/2020.01.07.898031

**Authors:** Yijie Wang, Justin M Fear, Isabelle Berger, Hangnoh Lee, Brian Oliver, Teresa M Przytycka

**Author notes:** co-first author (Yijie Wang, Justin M Fear).

## Abstract

Gene Regulatory Networks (GRNs) control many aspects of cellular processes including cell differentiation, maintenance of cell type specific states, signal transduction, and response to stress. Since GRNs provide information that is essential for understanding cell function, the inference of these networks is one of the key challenges in systems biology. Leading algorithms to reconstruct GRN utilize, in addition to gene expression data, prior knowledge such as Transcription Factor (TF) DNA binding motifs or results of DNA binding experiments. However, such prior knowledge is typically incomplete hence resulting in missing values. Therefore, the integration of such incomplete prior knowledge with gene expression to elucidate the underlying GRNs remains difficult.

To address this challenge we introduce NetREX-CF – Regulatory **Net**work **R**econstruction using **EX**pression and **C**ollaborative **F**iltering – a GRN reconstruction approach that brings together a modern machine learning strategy (Collaborative Filtering model) and a biologically justified model of gene expression (sparse Network Component Analysis based model). The Collaborative Filtering (CF) model is able to overcome the incompleteness of the prior knowledge and make edge recommends for building the GRN. Complementing CF, the sparse Network Component Analysis (NCA) model can use gene expression data to validate the recommended edges. Here we combine these two approaches using a novel data integration method and show that the new approach outperforms the currently leading GRN reconstruction methods.

Furthermore, our mathematical formalization of the model has lead to a complex optimization problem of a type that has not been attempted before. Specifically, the formulation contains *ℓ*_0_ norm that can not be separated from other variables. To fill this gap, we introduce here a new method Generalized PALM (GPALM) that allows us to solve a broad class of non-convex optimization problems and prove its convergence.

## 1 Introduction

Regulation of gene expression is central to cellular function. The regulatory relationships between transcription factors (TFs) and the genes they target (TGs) are captured by the Gene Regulatory Network (GRN). Inference of these cell type specific GRNs is a current challenge in systems biology. Earlier work focused on predicting regulatory networks using gene expression data alone, but these methods tend to have poor predictive power [1, 2]. Indeed, inference of network edges based solely on gene expression data is challenging; network reconstruction uses an enormous search space, and the underlying biology is multilayered with many factors including post-transcriptional and post-translation regulation contributing to TF’s activity. We and others have found that network accuracy is drastically improved by including additional biological data such as chromatin structure (i.e., ATAC-Seq and ChIP-Seq), TF DNA binding motifs, and DNA sequence conservation scores [3, 4, 5, 2, 6, 7].

Additional biological data have been used as *a prior* to inform network model selection in a variety of contexts [7, 8, 9]. MerlinP [5] uses network priors to influence the objective function for model selection. While, Inferelator [3], a method built on network component analysis (NCA), uses given gene expression data and a network prior to estimate TF activity. Furthermore, Inferelator predicts the GRN by uncovering the relationship between TF activity and their target genes’ expression. We recently developed NetREX [2], which is also based on the NCA model, but NetREX simultaneously estimates TF activity while modifying the prior network by adding and removing edges.

Because of the NCA model’s simplicity yet biological relevance, this approach becomes the foundation of the current state-of-the-art methods for GRN reconstruction [10, 11, 12, 3, 4, 2, 13, 14]. NCA uses the prior network’s structure to inform the decomposition of gene expression into TF activities [10]. Specifically, TF activities are modelled as a hidden variable accounting for the complex and often unknown relationships between TF expression and TF regulatory activity. TF activity is more robust and has been proved to be superior to TF gene expression in the task of GRN reconstruction [3]. However, NCA-based methods heavily rely on the quality of the prior network. If a prior network is very noisy, NCA-based methods cannot reliably predict TF activity, and in such circumstances the GRNs predicted by those methods are not trustworthy [2]. Therefore, building a reliable prior network becomes the key factor to employ the NCA-based methods.

A GRN prior is typically built by integrating various types of biological data, but construction of a quality prior is challenging due to the incompleteness of available data. For example, we can build a prior network by using TF-DNA binding data (e.g. ChIP-seq). However, we often only have access to ChIP-seq data for a fraction of TFs. Therefore, all interactions with TFs that do not have ChIP-seq data are considered as missing values. Similarly, computational mapping of TF-DNA binding motifs may miss true physical binding sites due to the problem of multiple testing, leading to incompleteness in the TF-DNA motif prior. Current methods for building GRNs by integrating multiple sources of prior knowledge do not directly account for the fact that there is missing data [7]. However, in the last decade we have witnessed a rapid development of machine learning methods capable dealing with large amounts of missing data. One particularly successful approach is Collaborative Filtering (CF), the method used by NETFLIX’s movie recommendation system [15, 16]. Given incomplete information about a user’s preferences, CF infers informative features and then applies them to provide movie recommendation for other users in the absence of complete information.

In this work, we present NetREX-CF – Regulatory **Net**work **R**construction using **EX**pression and **C**ollaborative **F**iltering – a GRN reconstruction approach that uses the idea of CF in a completely novel way, namely by combining such recommendation system with expression-based model optimization. Similar to its precursor, NetREX, NetREX-CF selects a network model by simultaneously optimizing network topology and its NCA-based fit of gene expression data. However, rather than arriving to a final network by reprogramming the edges in the prior network, NetREX-CF uses a joint optimization function to directly integrate expression data with other types of prior knowledge using CF. We demonstrate that CF takes the fullest advantage of the prior data, and when combined with the biologically relevant NCA-based model, provided a remarkable improvement over existing approaches.

Mathematically, the simultaneous optimization of network topology, fit of the NCA model, and feature selection for the CF yielded a complex optimization problem of a type that has not been attempted before. Specifically, the optimization is non-convex and non-smooth due to the binary nature of presence/absence of network edges. More importantly, the optimization contains *ℓ*_0_ norm that can not be separated from other variables that need to be optimized. While the recently introduced PALM method [17] can solve a certain class of such non-convex optimization problems, where the *ℓ*_0_ norm is separable (in particular the one used in NetREX), a simultaneous optimization of all three sets of parameters yields a problem that cannot be solved by PALM. To fill this gap, we introduce GPALM (Generalized PALM), a new provably convergent method for solving a broad class of non-convex optimization problems with an inseparable *ℓ*_0_ norm. Therefore, in addition to introducing a new method to reconstruct GRNs that outcompetes previous methods, this work also provides a solution to an important class of optimization problems.

## 2 NetREX-CF - Method Overview

The NetREX-CF model is a novel data integration framework for reconstructing GRNs by organically utilizing both gene expression *E* and a set of prior networks *P* = {*P*^1^, …*P*^*d*^}. The main idea behind the NetREX-CF model is an integration of two complementary optimization strategies: (i) a machine learning component designed based on Collaborative Filtering that is able to identify hidden features from the current observed prior networks *P* and utilize these features to recommend an improved GRN and (ii) a sparse NCA-based network remodelling component that can refine the topology of a GRN based on given gene expression *E*. These two computational components operate alternatively. The CF component recommends new edges to the current GRN and the sparse NCA-based network remodelling component screens the recommended edges and keeps the edges that are essential to explain the given gene expression. Once the sparse NCA-based network remodelling component confirms some of the recommended edges, the CF component further utilizes those retained recommended edges to make new edge recommendations for the sparse NCA-based network remodelling component to further examine (illustrated in Fig. 1).

**Fig. 1:**
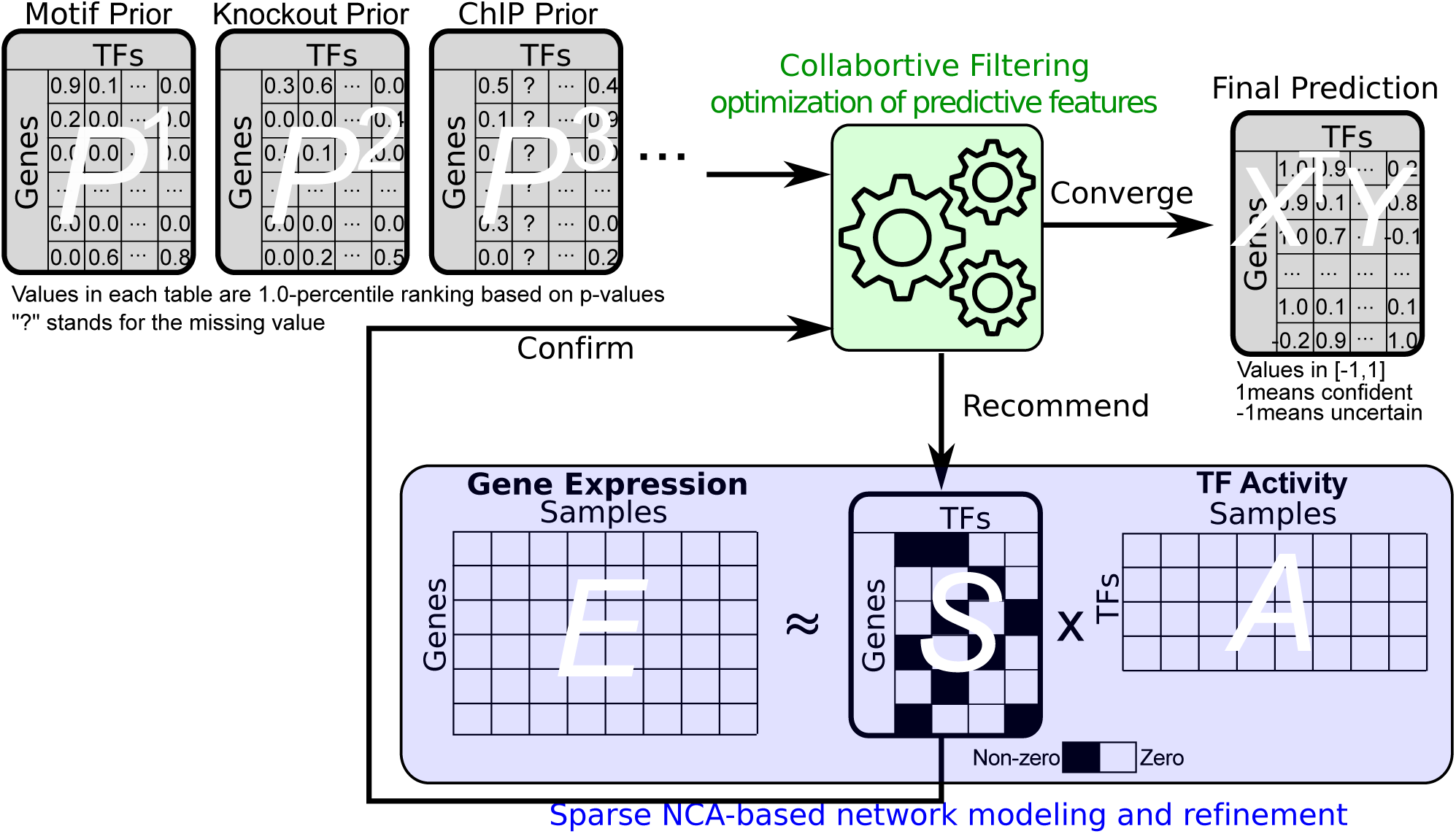
Method Overview. Collaborative Filtering (CF) and NCA-based gene expression modelling alternatively inform each other during a joint optimization process: CF recommends edges to be confirmed by the NCA model and used to improve CF.

Computationally, this is achieved by a simultaneous optimization of the following sets of variables: (i) the activities of TFs (matrix *A*), (ii) a weighted GRN (matrix *S*), and (iii) two feature matrices: the hidden features for target genes (*X* where the *i*th row *x*_*i*_ represents the hidden feature vector for gene *i*) and the hidden features for TFs (*Y* where the *j*th row *y*_*j*_ represents the hidden feature vector for TF *j*). The matrix *A* is optimized by the sparse NCA-based network remodelling component while the matrices *X* and *Y* are optimized by the Collaborative Filtering component. Notably, matrix *S* is the connection between the two components and should be optimized by considering both components.

Formally, *E* ∈ ℝ^*n*×*l*^ is the matrix of expression data of *n* genes in *l* experiments and prior network *P* ^*k*^ ∈ ℝ^*n*×*m*^, ∀*k* is a weighted adjacency matrix of the bipartite graph that records the prior knowledge of regulations between *m* TFs and *n* genes. Matrix *A* ∈ ℝ^*m*×*l*^ is the TF activity for *m* TFs in *l* samples and *S* ∈ ℝ^*n*×*m*^ is a weighted GRN. We further define penalty matrix *C* and observation matrix *B* based on the set of prior networks *P*. Each element in *C* can be computed by 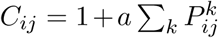 (*a* = 60 suggested by [16]) and each element in *B* is binary and can be computed by *B*_*ij*_ = 1 if 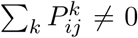 and *B*_*ij*_ = 0 otherwise. *X* ∈ ℝ^*n*×*h*^ contains feature vector *x*_*i*_ for gene *i* and *Y* ∈ ℝ^*m*×*h*^ contains feature vector *y*_*j*_ for TF *j*. Then, our optimization problem is formalized as following:

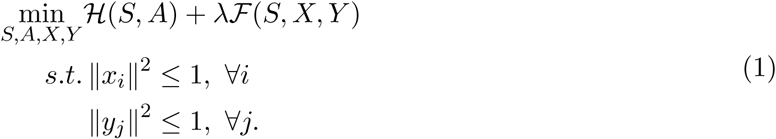

where:

– 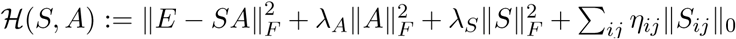 is the sparse NCA-based network remodelling component; 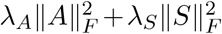 are standard regularization terms and Σ_*ij*_ *η*_*ij*_ ‖*S*_*ij*_ ‖_0_ induces sparsity of a given prior GRN and therefore only essential edges that help to minimize ℋ(*S, A*) are retained. ‖*S*_*ij*_‖_0_ is the *ℓ*_0_ norm that is 1 if *S*_*ij*_ ≠ 0 and 0 otherwise.
– 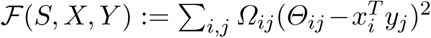 optimizes the hidden features *X* and *Y* of the Collaborative Filtering component; *Θ*_*ij*_ is a binary matrix of edges to be predicted by the hidden features in the given iteration and *Ω*_*ij*_ encodes penalties that guide the predictions. Both *Θ*_*ij*_ := ‖*S*_*ij*_‖_0_⊕*B*_*ij*_ and 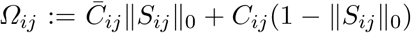 are defined based on ‖*S*_*ij*_‖_0_ and the prior information (*C*_*ij*_). Detailed explanation of *Θ*_*ij*_ and *Ω*_*ij*_ are provided in Method Details section. For the initialization step, both *Θ*_*ij*_ and *Ω*_*ij*_ are defined based on the prior networks only while in the subsequent steps they also take into account the results of the sparse NCA-based network remodelling component (illustrated in Fig. 1 and Fig. 3).

To solve problem (1), we first put all continuous terms together and define 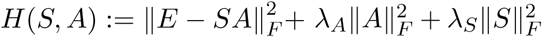 and non-continuous terms together and define 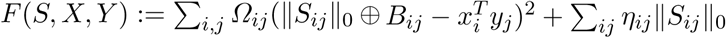. Then the optimization problem has a general format of an objective function as *Φ*(*S, A, X, Y*) = *H*(*S, A*)+ *F*(*S, X, Y*), where *H*(*S, A*) is continuous but non-convex and *F*(*S, X, Y*) is a composite function of *ℓ*_0_ norm of elements of *S* and other variables so it is neither continuous nor convex. More importantly, ‖*S*_*ij*_‖_0_ is coupled with *x*_*i*_ and *y*_*j*_, so that ‖*S*_*ij*_‖_0_ can not be separated from *F*(*S, X, Y*) as a separated term. To the best of our knowledge, there has been no known method that can optimize such a complex and non-convex function involving inseparable *ℓ*_0_ norm. To fill this gap, we propose here a new algorithm, Generalized PALM (GPALM) that generalizes the so called PALM method [17] and solves a class of problems of the format above, under the assumption that *F*(*S, X, Y*) is lower semi-continuous (see Supplementary Material A). In the Supplementary Material B, we propose the new GPALM method and prove its convergence.

## 3 Experimental Results

To demonstrate the capability of our proposed GRN reconstruction method, we collect multiple datasets that measure different perspectives of the gene regulation in yeast. These datasets include TF ChIP [5, 18, 19], TF DNA binding motif [5, 20], genetic knockout [5, 21, 22], and yeast gene expression [5, 23, 24, 25]. TF ChIP, motif, and genetic knockout datasets serve as prior knowledge for TF-gene interactions in the yeast GRN. The details of these priors are summarized in Table 1 and the overlap among priors is illustrated in Table 1. We further utilize TF-gene interactions extracted from YEASTRACT database [26] as a gold standard to validate the performance of GRN reconstruction. These gold standard TF-gene interactions are supported by both DNA binding and expression evidence. The details of the gold standard TF-gene interactions and their overlap with the prior datasets are shown in Table 1. Results generated by NetREX-CF are benchmarked against the results obtained from the published sequential methods. In the following, we detail the comparison between NetREX-CF, MerlinP [5], NetREX [2], LassoStARS [4], the original CF [16], and the summation of all prior knowledge (PriorSum). For a detailed description of parameter selection for competing methods, we refer the reader to the Supplementary Material D.

**Table 1:** Overlap between prior networks and the gold standard network.

| Network | # Genes | # TFs | # Edges | #Overlap<br>with Motif | #Overlap<br>with Knockout | #Overlap<br>with ChIP | #Overlap<br>with YEASTRACT |
| --- | --- | --- | --- | --- | --- | --- | --- |
| Motif | 5,506 | 197 | 187,079 | 187,079 (100%) | 9,236 (8.4%) | 8,717 (3.5%) | 3,497 (31.0%) |
| Knockout | 5,543 | 262 | 96,809 | 9,236 (4.6%) | 96,809 (100%) | 7,027 (2.9%) | 3,050 (27.0%) |
| ChIP | 5,557 | 318 | 229,936 | 8,717 (4.3%) | 7,027 (6.4%) | 229,936 (100%) | 2,656 (23.5%) |
| YEASTRACT | 3,731 | 148 | 10,525 | 3,497 (1.7%) | 3050 (2.8%) | 2,656 (1.1%) | 10,525 (100%) |

The last four columns show overlap between different networks and parentheses shows the corresponding percentage. YEASTRACT is the gold standard network we used for performance comparison.

To ensure an impartial comparison, we use average percentile ranking scores. For each method and for each gene *i*, we can obtain a list of TFs that are predicted to regulate gene *i* and sort these TFs by the confidence of the prediction (most confident at the top). We use 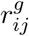 to denote the percentile-ranking of TF *j* within the ordered list of all TFs for gene *i*. Thus, 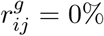 means that TF *j* is predicted with the highest confidence to regulate gene *i*, preceding all other TFs in the list. Based on the gold standard TF-gene interaction dataset *I*, we set *I*_*ij*_ = 1 if TF *j* regulates gene *i* in the gold standard dataset and *I*_*ij*_ = 0 otherwise. For any gene *i*, we use the average rank of the gold standard edges in the list of TF predicted to regulate gene *i* as the measure quality of the prediction:

**Fig. 2:**
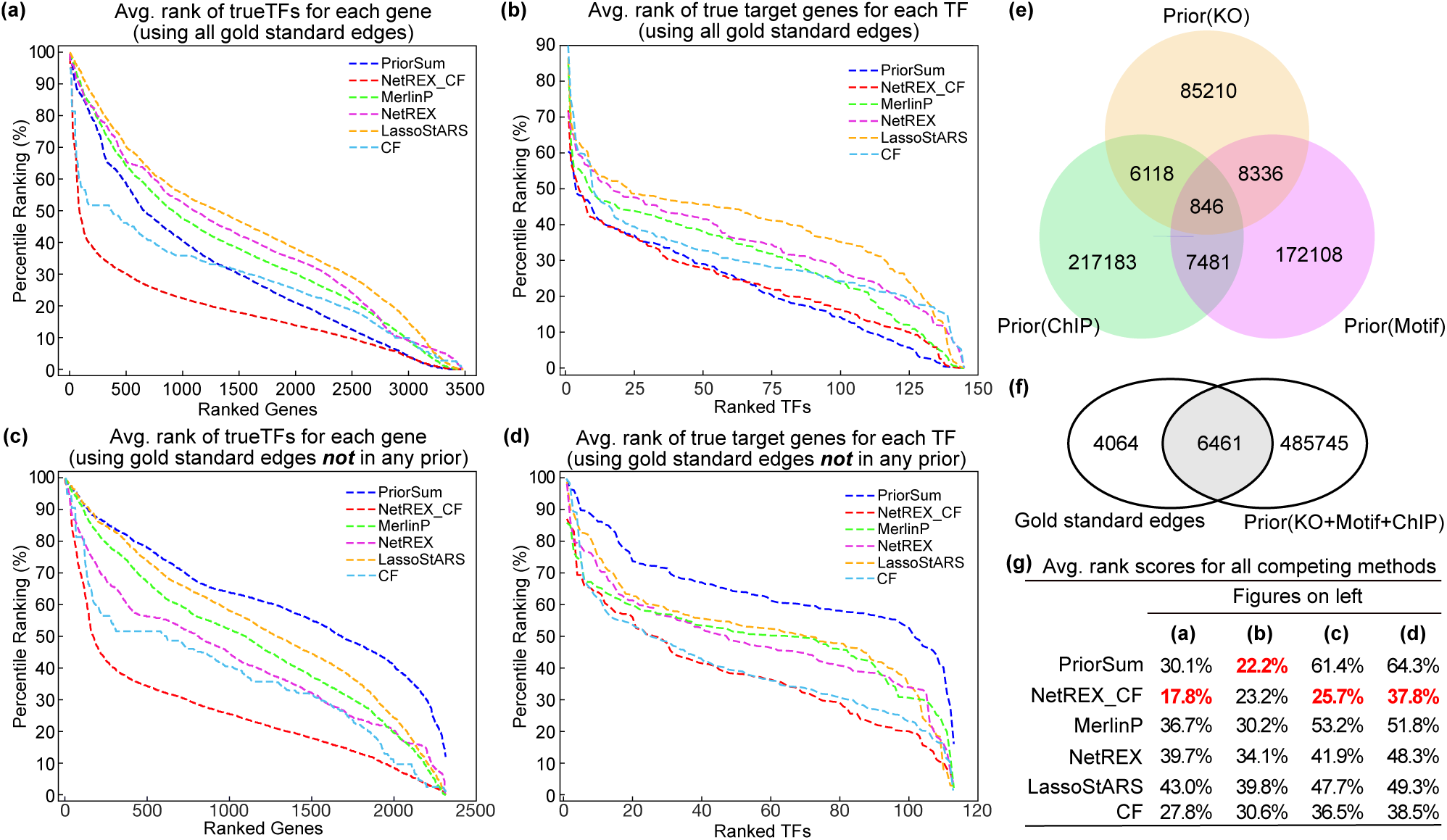
Performance comparison for all competing algorithms on the yeast dataset. **(a)** The performance of the methods on the task of predicting regulating TFs: for each algorithm, we compute for each gene the rankings of gold standard edges 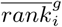 adjacent to it, sort them in descending order, and plot the sorted average rankings. **(b)** The performance of the methods on the task of predicting regulated genes using a measure similar as in (a) but focusing on genes regulated by TFs. **(c)** The performance of the methods on the task of predicting regulating TFs that are not observed in prior data. The procedure is the same as in (a) but only the gold standard edges that are not included in the prior knowledge are used for the evaluation. **(d)** The performance of the methods on the task of predicting regulated genes not observed in prior data (similar to (c) but focusing on genes regulated by TFs). **(e)** Overlap between priors. **(f)** Venn diagram for gold standard dataset and the union of the three prior datasets. **(g)** Summary of the average rankings of each algorithm for tasks reported in panels (a), (b), (c), and (d).

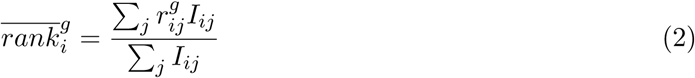

Lower values of 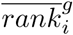 are more preferable, as they indicate gold standard TFs for gene *i* have lower rank than others. Furthermore, the over all ranking considering all genes can be computed by

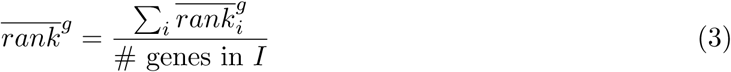

The denominator is the number of genes that have gold standard TFs in dataset *I*.

Similarly, for each TF we can measure the quality of the sorted list of genes predicted to be regulated by it:

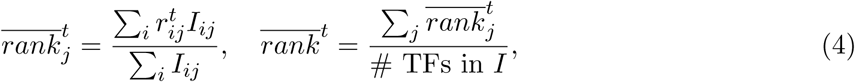

where 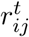 denotes the percentile-ranking of gene *i* with in the ordered list of all genes for TF *j* and 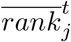 is the average rankings for the gold standard genes for TF *j*. 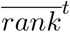 is the overall average rankings considering all TFs.

Fig. 2 (a) illustrates the comparison between the competing algorithms in terms of average rankings of gold standard TFs for each target gene. As shown, the sorted average ranking curve for NetREX-CF is below all other methods, indicating that the average rankings of gold standard TFs predicted by NetREX-CF for each gene are much lower than the rankings predicted by other methods. In the average rankings of gold standard genes among the genes predicted to be regulated by each TF, surprisingly, PriorSum (the weighted edge summation of three priors) outperforms all previous computational methods by a large margin. In contrast, NetREX-CF is competitive with PriorSum (Fig. 2 (b)) indicating that it takes the best advantage of the prior data. Notably, NetREX-CF outperforms the original CF, which demonstrates that the integration of CF model and sparse NCA-based model is beneficial.

Next, in order to demonstrate the advantages of NetREX-CF in predicting ranks for missing data (edges that does not appear in the prior knowledge datasets), we identified all edges that are in gold standard dataset but are not supported by any prior dataset. Indeed, as shown in Fig. 2 (f), a large portion of gold standard dataset (4,064 out of 10,525 gold standard TF-gene interactions) are not covered by any prior dataset. Therefore, we can use these gold standard interactions with missing prior data to compare the ability of the competing methods in recovering rankings under the assumption of missing data. As shown in Fig. 2 (c) and (d), NetREX-CF achieves much lower rankings for those missing data. The curves of NetREX-CF in Fig. 2 (c) and (d) are below curves of other methods by large margins except for Fig. 2 (d), where NetREX-CF is marginally better than the original CF demonstrating the benefits of integrating the CF method for predicting target genes. As shown in Fig. 2 (g), NetREX-CF achieves the lowest overall average ranking scores for all but one task where its performance is competitive with the winning method.

## 4 Method Details

We now describe our method in more detail. We first elucidate the NetREX-CF model presented in (1). Then, we illuminate the specified GPALM algorithm we developed to solve the NetREX-CF model.

### 4.1 NetREX-CF Model

Before describing the mathematical foundation of the NetREX-CF model, we provide a brief overview of Collaborative Filtering model and the sparse NCA-based network remodelling model, respectively. Next we formally introduce the integration of these two models.

#### Collaborative Filtering Model

As illustrated in Fig. 3, to reconstruct GRNs we might have access of several prior networks, each of which reflects different perspective of the gene regulation process. Here we illustrate three prior networks: the Motif prior network, the Knockout prior network, and the ChIP prior network. In general, the prior networks are partial observation of the gene regulation process and therefore incomplete. The incompleteness of prior networks can be further demonstrated by Table 1, where there are only a small number of overlaps between the yeast prior networks and the gold standard GRN. Previous prior-based GRN reconstruction methods [7] typically make efforts to preserve those edges in the prior networks into the final GRN reconstruction but are unable to predict new edges to resolve the incompleteness of the prior networks.

**Fig. 3:**
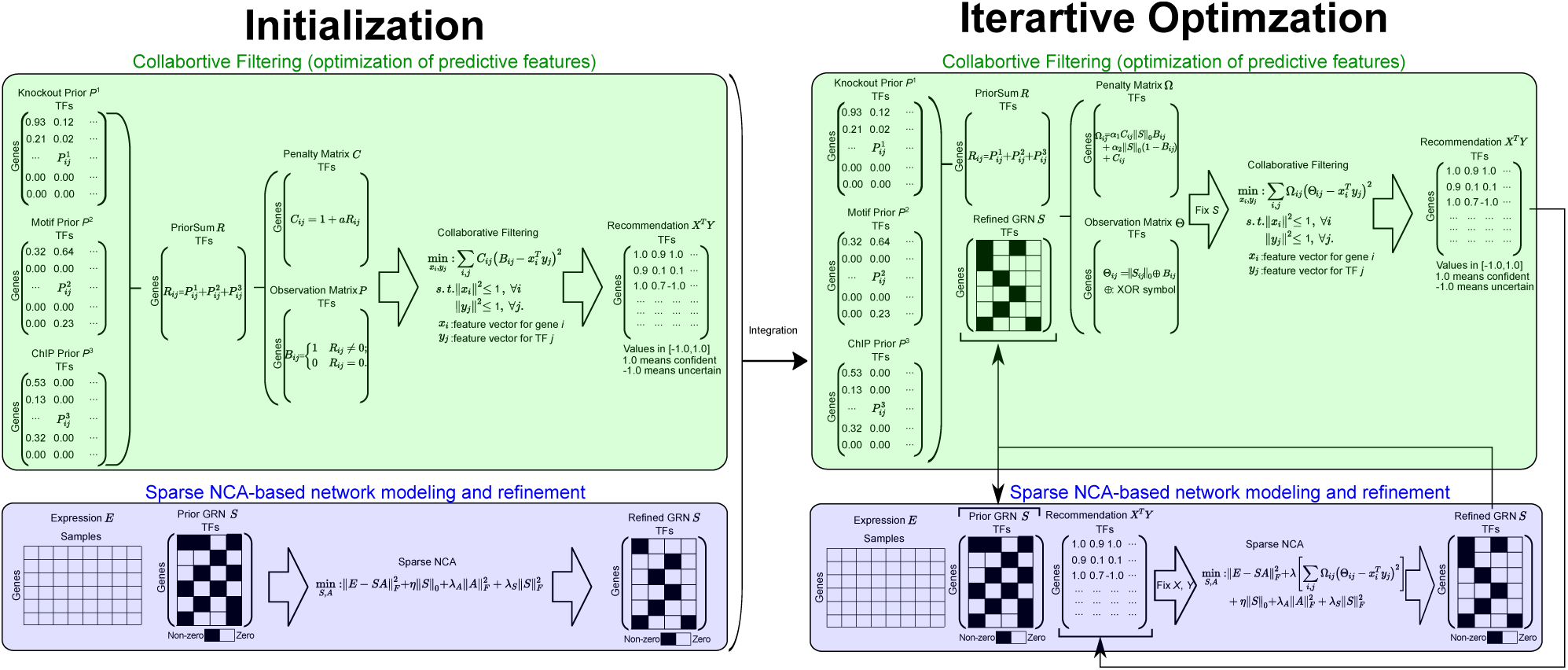
The overview of the information flow in the NetREX-CF optimization.

Collaborative filtering, a machine learning technique, is an approach to mitigate the incompleteness of the prior networks. Collaborative filtering is able to make prediction based on partial observation. Given a set of prior networks *P* = {*P*^1^, …*P*^*d*^}, the mathematical formulation of collaborative filtering can be presented by

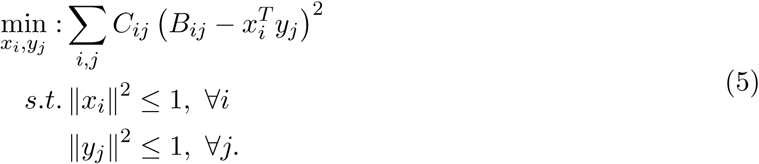

We recall that *x*_*i*_ and *y*_*j*_ are hidden feature vectors for gene *i* and TF *j*, respectively, and *B*_*ij*_ is a binary number that equals to 1 when we observe the edge between gene *i* and TF *j* in any prior and equals to 0 otherwise. *B*_*ij*_ encodes that predictions that feature vectors need to make. 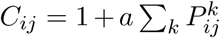 is the penalty for learning the edge between gene *i* and TF *j*. Larger *C*_*ij*_ implies *B*_*ij*_ = 1 and also encourages the dot product 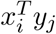 between gene feature vector *x*_*i*_ and TF feature vector *y*_*j*_ to be 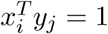. Details of the CF model is illustrated in Fig. 3 top left panel.

After solving the optimization problem (5), we can use 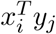, ∀*i, j* to predict edges that are not in the prior networks. Because of the constraints in (5), we know 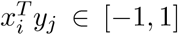 based on Cauchy–Schwarz inequality. 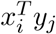 is close to 1 implies that the collaborative filtering method recommends the edge between gene *i* and TF *j*. However, to obtain reliable predictions, it is beneficial that the correctness of the edge recommendation is further confirmed by other methods.

#### Sparse NCA-based Network Remodelling Model

Other than utilizing prior information, such as binding properties, we can use gene expression to help build reliable GRNs. Currently, the state-of-art methods to use gene expression for reconstructing GRNs are NCA-based approaches [10, 11, 12, 3, 4, 2, 13, 14]. However, in order to use the NCA model, we need a prior network in addition to gene expression data. Given gene expression *E* ∈ ℝ^*n*×*l*^ for *n* genes in *l* samples and a prior network *S* ∈ ℝ^*n*×*m*^, the sparse NCA-based network remodelling model can be presented as

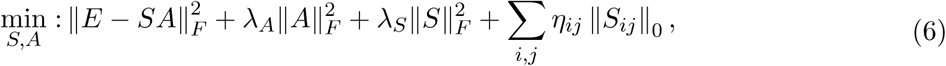

where the first term is the basic NCA model [10] (*A* ∈ ℝ^*m*×*l*^ is the TF activity for *m* TFs in *l* samples) and the second and third terms are standard regularization terms and the last term involving *ℓ*_0_ norm that is able to induce sparsity of the given prior network. Therefore, solving (6) would yield a refined GRN that only retains key edges from the prior network. The details of the sparse NCA-based network remodelling model is illustrated in Fig. 3 bottom left panel. Since for most of cases, we do not have a prior network, we need to build a reliable prior based on multiple sources of prior information.

#### Formulation of the NetREX-CF Model

Here we propose to integrate both CF model and Sparse NCA-based Network Remodelling model. As we mentioned that the CF model needs a way to confirm the recommended edges and the sparse NCA-based network remodelling model needs a prior network to work with. Therefore, it is very natural to combine theses two models together. The CF model can recommend a prior network for the sparse NCA-based network remodelling model, and as a reward, the sparse NCA-based network remodelling model is able to confirm the recommend edges and thus allow the CF model to predict new edges. The mathematical formulation of the NetREX-CF model is

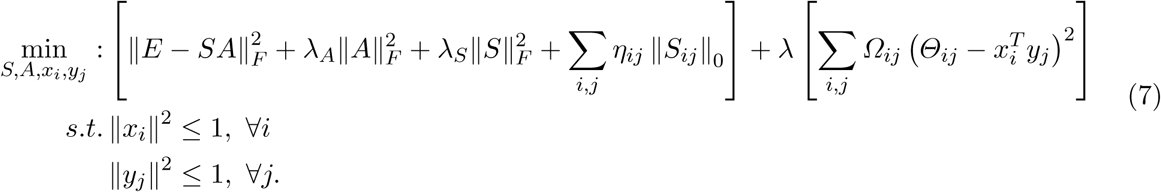

The first square bracket is the sparse NCA-based model and the second square bracket is the CF model. *λ* is the balance between these two models. In the NetREX-CF model, we define *Θ*_*ij*_ = ‖*S*_*ij*_ ‖_0_ ⊕ *B*_*ij*_ = ‖*S*_*ij*_‖_0_ + (1 − ‖*S*_*ij*_‖_0_)*B*_*ij*_ (⊕ is XOR operation) to let the CF model not only predict edges in the prior networks *B*_*ij*_, but also take into account the edges confirmed by the sparse NCA-based model *S*_*ij*_. Furthermore, *Ω*_*ij*_ is defined as 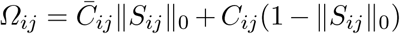, where 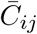 is the user defined penalty for edges confirmed by the sparse NCA-based model *S*_*ij*_ ≠ 0 and *C*_*ij*_ is the penalty for edges not in *S* (*S*_*ij*_ = 0). The details of the NetREX-CF model is illustrated in Fig. 3 right panel. The details of how to select all the user defined parameters of the NetREX-CF model are elaborated in the Supplementary Material D.

Once we put the definition of *Ω*_*ij*_ and *Θ*_*ij*_ into (7) and we put Σ_*ij*_ *η*_*ij*_‖*S*_*ij*_‖_0_ into the second square bracket, we have

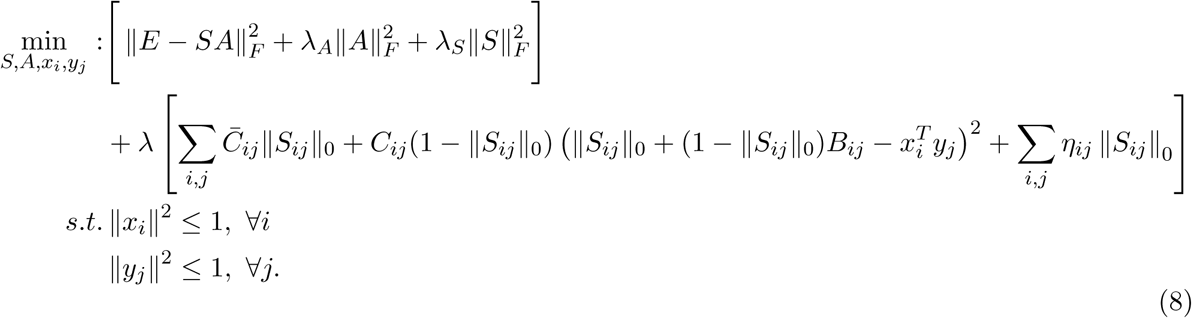

Then the function in the first square bracket is continuous and we define it as *H*(*S, A*). The function in the second square bracket is lower semi-continuous (Supplementary Material D.6) and we define it as *F*(*S, X, Y*). Clearly, we cannot separate ‖*S*_*ij*_‖_0_ from *x*_*i*_ and *y*_*i*_ and put every term involving ‖*S*_*ij*_‖_0_ together as a separated term. To the best of our knowledge, there is no known method that is able to solve the optimization problem (8). In the following, we elaborate the algorithm we developed to solve the NetREX-CF model.

### 4.2 The NetREX-CF Algorithm

Because current methods can not solve problem (8), we propose a Generalized PALM (GPALM) algorithm that is an extension of the PALM algorithm [17]. GPALM can be used to solve this class of optimization problem involving inseparable *ℓ*_0_ norm, which is when *ℓ*_0_ norm cannot be separated from other optimized variables as a separated term. The format of the problem that GPALM can solve is provide in the Supplementary Material A. The GPALM algorithm and its convergence proof are provided in the Supplementary Material B. Here we directly applied the GPALM algorithm to solve our NetREX-CF model. The algorithm is listed as follows. The proximal operator used in the algorithm is defined as:

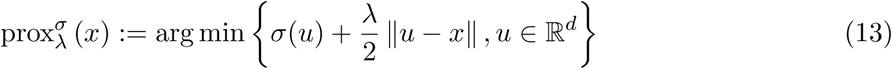

#### Algorithm 1: The algorithm for problem (M).

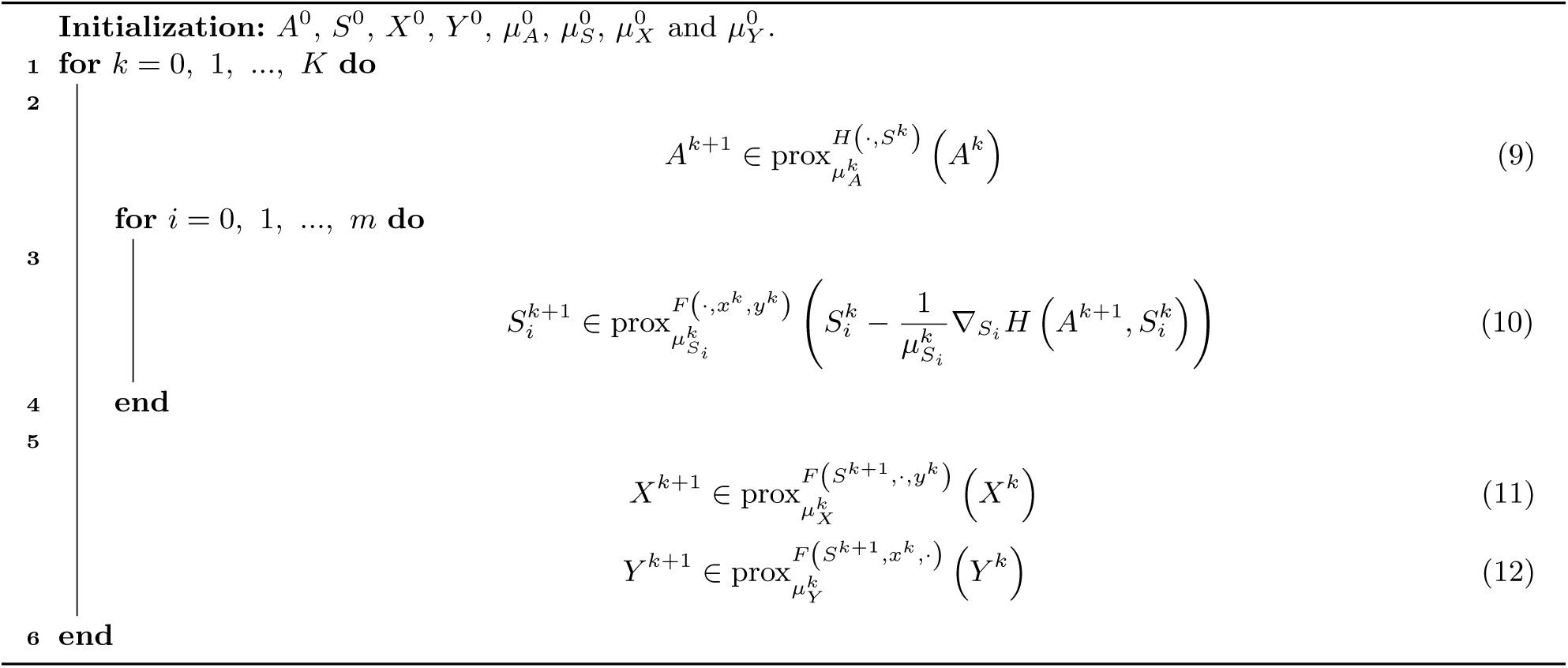

The proximal operator and proximal gradient methods are often applied to replace conventional smooth optimization techniques for functions that are not continuous but can be approximated by well behaving functions (or have other nice bounding properties).

We show in the following that, for all proximal operators used in the above algorithm, we can compute the corresponding update steps by either using a closed form that we are able to derive or by reducing the computation to a convex optimization problem.

**Update *A*** The proximal operator (9) has a closed form solution.

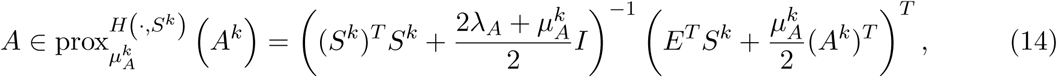

where 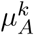 is the Lipschitz constant that can be computed by 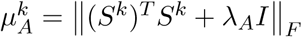. The details of the derivation related to update *A* can be found in Supplementary Material C.1.

**Update *S*** Similarly, the proximal operator (10) also has a closed form solution.

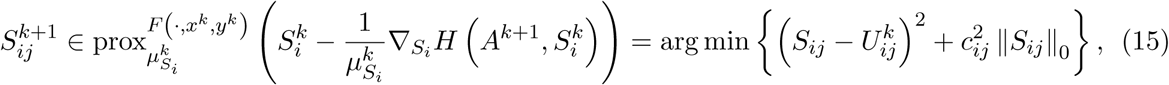

where 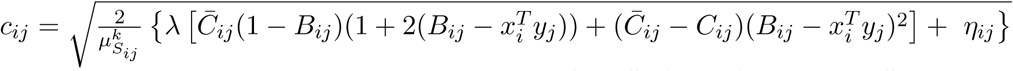 and 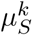 is the Lipschitz constant that can be computed by 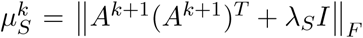. Therefore, the closed solution of the above problem is

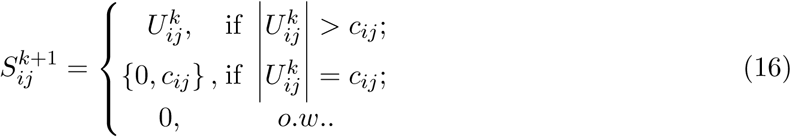

The details of the derivation related to update *S* can be found in the Supplementary Material C.2.

**Update *X*** Each row *x*_*i*_ of *X* needs to be updated by solving the following proximal operator.

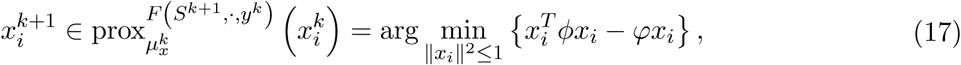

where 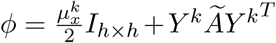 and 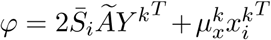. *Ã* be the diagonal matrix with the values *Ā*_*i*1_, *Ā*_*i*2_, ..*Ā*_*im*_ on the diagonal, where 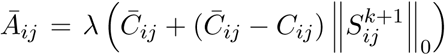. And 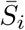 is defined as 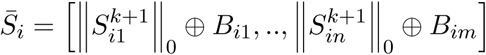. Since the problem becomes a Quadratically Constrained Quadratic Program (QCQP), we leave the rest to the CVXPY python package [27, 28]. The details of the derivation related to update *X* can be found in the Supplementary Material C.3.

**Update *Y*** Each row *y*_*i*_ of *Y* needs to be updated by solving the following proximal operator.

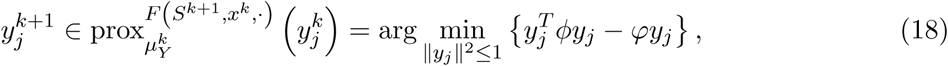

where 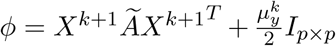 and 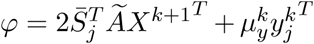. *Ã* that is also a diagonal matrix with the values *Ā*_1*j*_, *Ā*_2*j*_, ..*Ā*_*mj*_ on the diagonal and 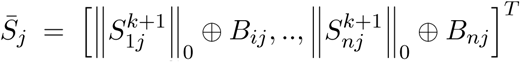. Since the problem also becomes a QCQP, we leave the rest to the CVXPY python package. The details of the derivation related to update *Y* can be found in the Supplementary Material C.4.

## 5 Conclusions

Data integration and predictive modelling are the two key tasks of Computational Biology. However, these two tasks are rarely considered together. GRN reconstruction is an example of an important and challenging computational biological problem that can benefit from both approaches. Here we propose a method that combines machine learning based data integration strategy and a gene expression modelling approach into one global iterative optimization strategy where machine learning component informs the expression based modeling component and vice versa. Our new integrative GRN reconstruction method outperforms previous computational methods for this task demonstrating the power of our integrative approach.

We believe that the general approach presented in this study provides not only an important step towards reconstructing better GRNs, but it has also a potential to become a paradigm for addressing other optimization problems in computational biology.

## Acknowledgement

This research was supported by the Intramural Research Program of the National Library of Medicine and the National Institute of Diabetes and Digestive and Kidney Diseases at the National Institutes of Health, USA.

## Supplementary Materials

We extend the original PALM algorithm [17] and propose the GPALM algorithm that can solve more general problems. The format of the problem that GPALM can solve is explained in section A. The actual algorithm and the its convergence proof are provided in section B.

### A GPALM Preliminary

#### A.1 The problem and basic assumptions

We are interested in solving the non-convex and non-smooth minimization problem with the following structure

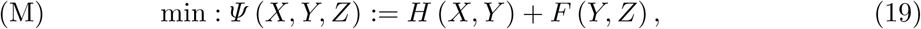

where we have the following assumption:

##### Assumption 1. The assumptions for problem (M) is as follow

1. *H* : ℝ^*n*^ × ℝ^*m*^ → ℝ *is a C*^1^ *function.*
2. *F* : ℝ^*m*^ × ℝ^*l*^ → (−∞ + ∞] *is a proper and lower semicontinuous (PLS) function. And F* (*Y, Z*) *has the following structure* 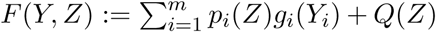, *where p*_*i*_ : ℝ^*l*^ → ℝ *is Lipschitz continuous with moduli L*_*i*_(*Z*) *and p*_*i*_(*Z*) > 0, ∀*i and g*_*i*_ : ℝ → ℝ *is lower semicontinuous and* sup *g*_*i*_(*Y*_*i*_) *< λ*_*i*_, ∀*i and Q* : ℝ^*l*^ → ℝ *is Lipschitz continuous with moduli L*_*Q*_(*Z*). *Y* = [*Y*_1_, *…, Y*_*i*_, *…, Y*_*m*_].
3. 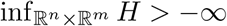 *and* 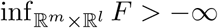.
4. *For any Y the function X* → *H*(*X, Y*) *is* 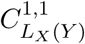, *namely the partial gradient* ∇_*X*_ *H*(*X, Y*) *is globally Lipschitz with moduli L*_1_(*Y*), *that is*

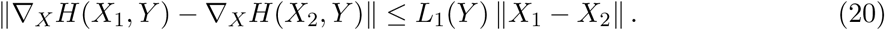 *Likewise, for any fixed X the function Y*_*i*_ → *H*(*X, Y*_*i*_) *is assumed to be* 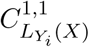.
5. *For any fixed Y the function Z* → *F*(*Y, Z*) *is assumed to be* 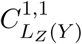.
6. ∇*H is Lipschitz continuous on bounded subsets of* ℝ^*n*^ × ℝ^*m*^. *In other words, for each bounded subsets T*_1_ × *T*_2_ *of* ℝ^*n*^ × ℝ^*m*^ *there exist M* > 0 *such that any* (*X*_1_, *Y*_1_) *and* (*X*_2_, *Y*_2_):

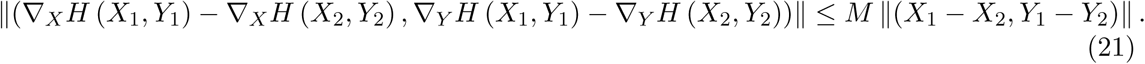

#### A.2 Subdifferentials of nonconvex and nonsmooth functions

##### Definition 1.

*Let σ* : ℝ^*d*^ → (−∞, +∞] *be a PLS function. For a given x* ∈ dom *σ, the Frechet subdifferential of σ at x, written* 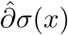, *is the set of all vectors u* ∈ ℝ^*d*^ *which satisfy*

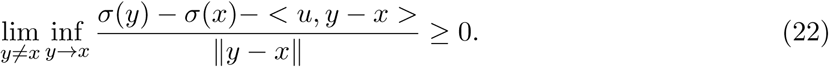

*When x* ∈ dom *σ, we set* 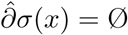.

##### Proposition 1.

*∂*(*λf* (*x*)) = *λ∂f* (*x*) *for any λ* > 0.

The proposition can be proved based Definition 1.

#### A.3 Proximal map

Let *σ* : ℝ^*d*^ → (−∞, +∞] be a PLS function. Given *x* ∈ ℝ^*d*^ and *t* > 0, the proximal map associate to *σ* id defined by:

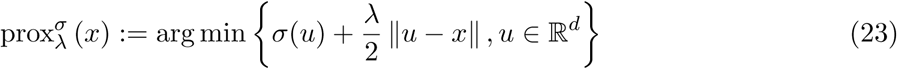

The proximal map has the following important property (Lemma 3.2 in []).

##### Lemma 1.

*Let h* : ℝ^*d*^ → ℝ *be a continuously differentiable function with gradient* ∇*h assumed L*_*h*_ *Lipschitz continuous and let σ* : ℝ^*d*^ → (−∞, +∞] *be a proper and lower semicontinuous function with* 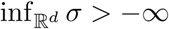. *Fix any t* > *L*_*h*_, *then for any u* ∈ dom*σ and any u*^+^ ∈ ℝ^*d*^ *defined by*

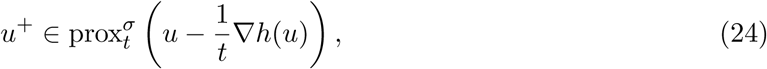

*we have*

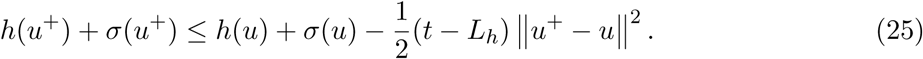

### B GPALM Algorithm and its Convergence Analysis

#### B.1 The Algorithm

Here we first write out the algorithm that is able to solve problem (M) with convergence guarantee.

##### Algorithm 2: The algorithm for problem (M).

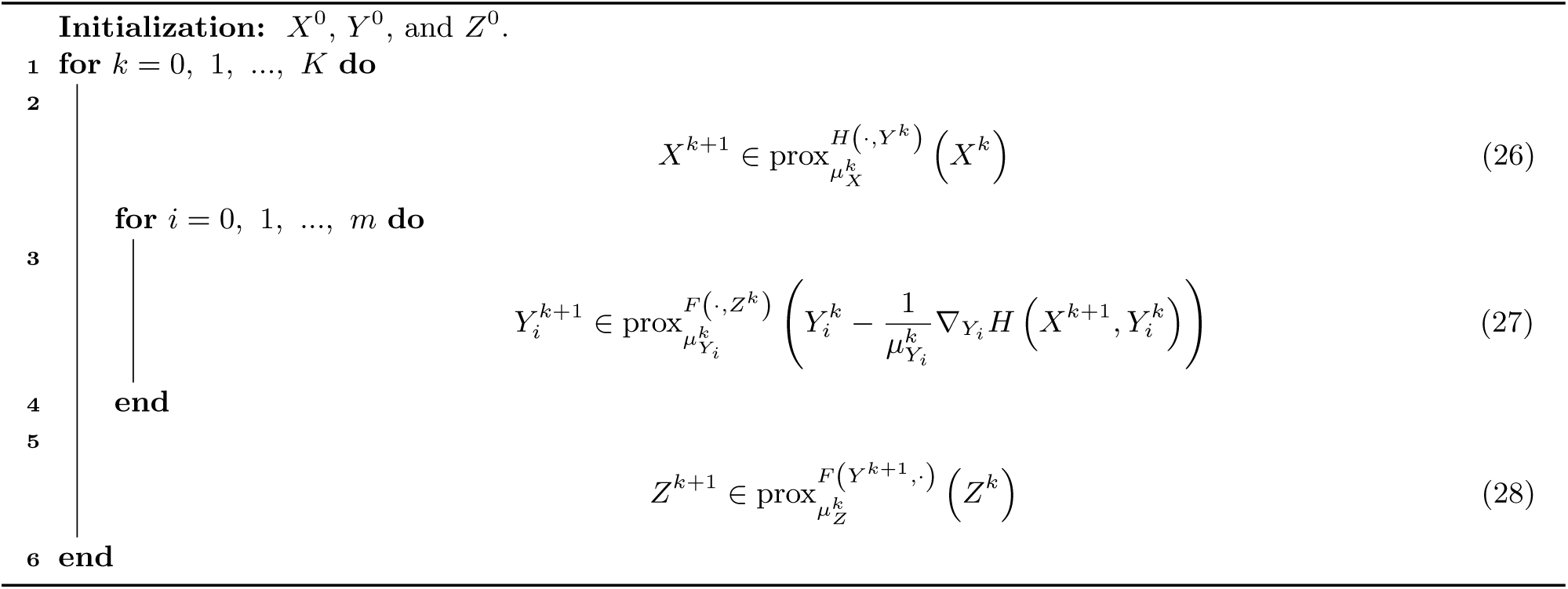

#### B.2 Convergence analysis

The proof procedure is followed the proofs introduced in the original PALM algorithm [17].

##### Theorem 1.

*Assume Ψ* (*B*) *is a PLS function with* inf *Ψ* > −∞, *the sequence* {*B*^*k*^}_*k*∈ℕ_ *is a Cauchy sequence and converges to a critical point of Ψ* (*B*), *if the following four conditions hold []:*

i. *Sufficiently decreasing: there exist some positive constant ρ*_1_ > 0, *such that*

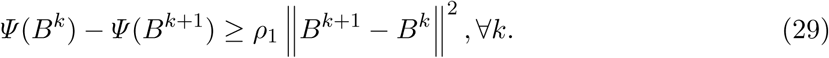
ii. *Relative error: there exist some positive constant ρ*_2_ > 0, *such that for any w*^*k*^ ∈ *∂Ψ* (*B*^*k*^),

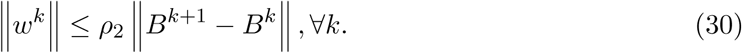
iii. *Continuity: there exist a subsequence*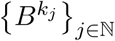 *and B*\*, *such that*

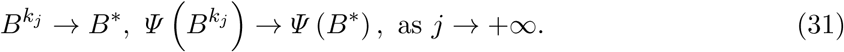
iv. *KL property: Ψ satisfies KL property in its effective domain.*

By the theorem above, we only need to check that the sequence generated by Algorithm 2 satisfy the conditions (i) - (iv).

##### Proposition 2. Algorithm 2 is a global convergence algorithm.

*Proof.* Follow Theorem 1, we prove Algorithm 2 satisfies conditions (i)-(iv).

**Condition (i)**. Based on (26), we know

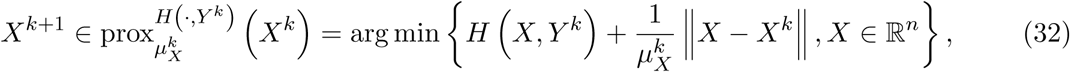

which implies

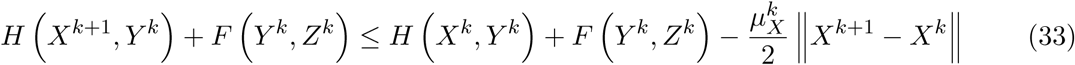

We then apply Lemma 1 to (27),

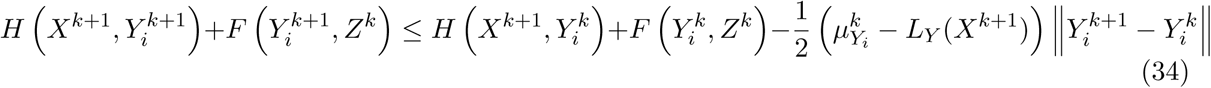

Similar to the derivation related to *X*, for *Z* we get

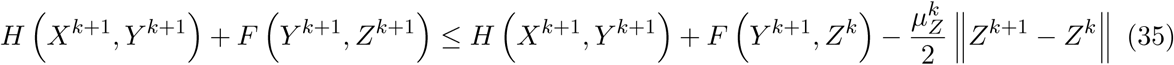

Let *B*^*k*^ = (*X*^*k*^, *Y* ^*k*^, *Z*^*k*^) and sum over equations from (33) to (35). We have

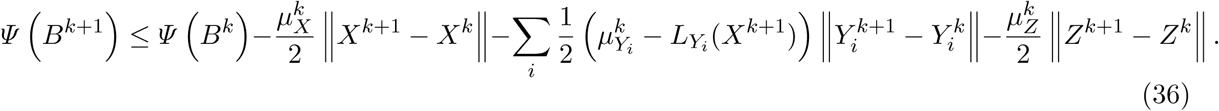

We know that 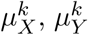, and 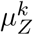 have their lower bound and 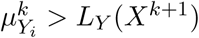. Therefore, we can get 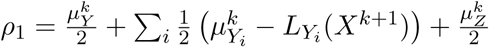. Then for *B*^*k*^ = (*X*^*k*^, *Y* ^*k*^, *Z*^*k*^) we have

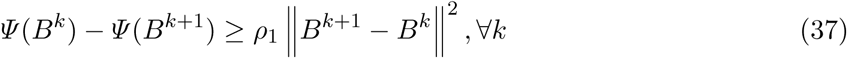

tha proves condition (i).

**Condition (ii)**. Writing down the optimality condition for (26), we have

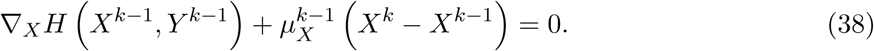

Let 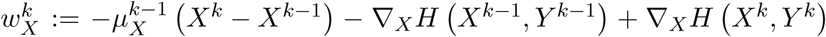. It is easy to prove that 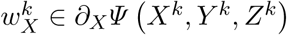. Then

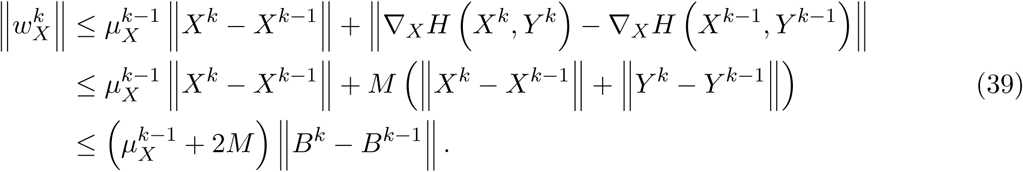

The first inequality comes from the fact that ∇*H* is Lipschitz continuous on bounded subset ℝ^*n*^ ×R^*m*^ as assumed in Assumption 1 (6).

The optimality condition for (27), we have

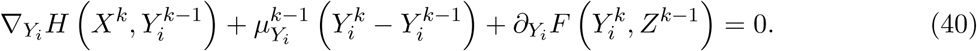

Let 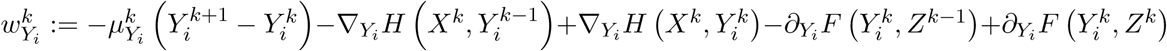. Clearly, 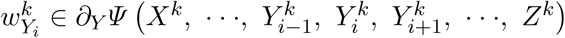, then we have

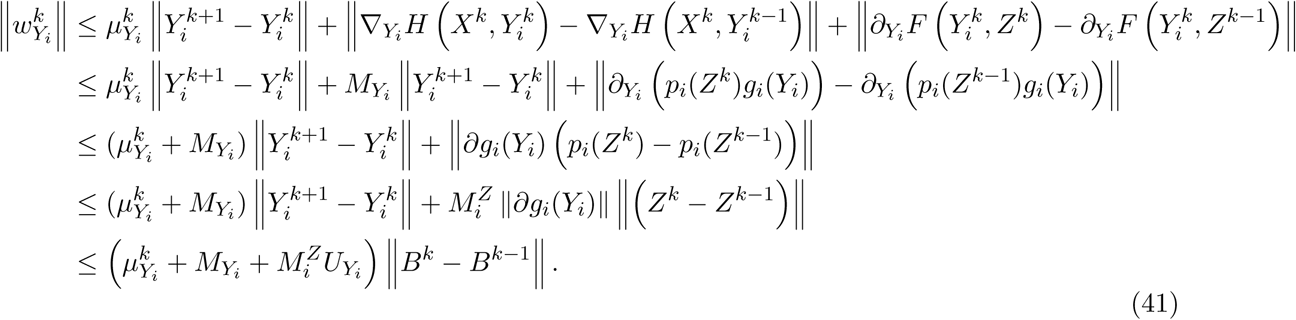

The second inequality utilizes the structure of *F*(*Y, X*) introduced in Assumption 1 (2). The third inequality uses Proposition 1. We set 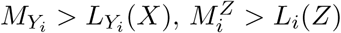, and 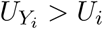.

Similar to things related to *X*, writing down the optimality condition for (28),

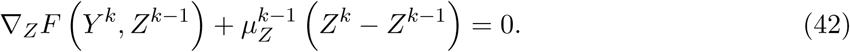

Let 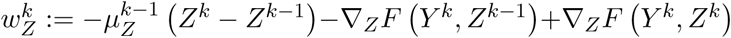. We find that 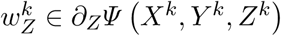 and we have

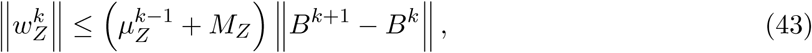

where *M*_*Z*_ > *L*_*Z*_ (*Y*).

Let 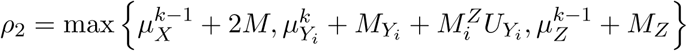 and sum (39), (41), (43), we have

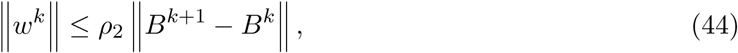

where 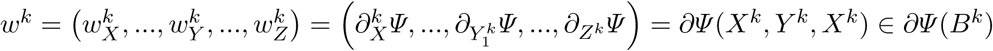.

**Condition (iii)**. Summing (37) from *k* = 0 to *N* − 1 we have

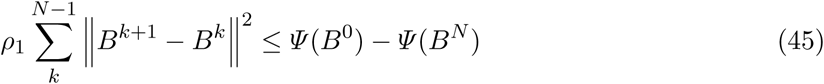

Since {*Ψ* (*B*^*N*^)} is decreasing and inf *Ψ* > −∞, there exist some 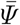 such that 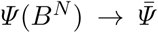 as. Therefore, let *N* → +∞ in (45), we have

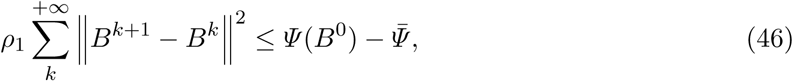

which implies that lim ‖*B*^*k*^ − *B*^*k*−1^‖ = 0. Let *B*\* = (*X*\*, *Y* *, *Z*\*) be a limit point of {*B*^*k*^}_*k*∈ ℕ_. = {(*X*^*k*^, *Y* ^*k*^, *Z*^*k*^)}_*k*∈ ℕ_. Then (46) indicates that there is a subsequence 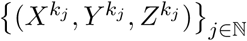 such that 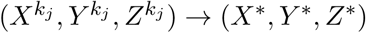 as *j* → + ∞.

From (27), we know

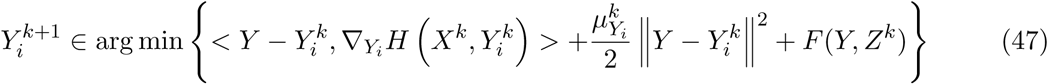

Let 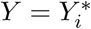 the limiting point of 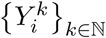, we have

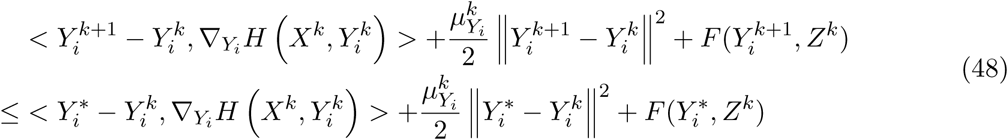

Set *k* = *k*_*j*_ − 1, we obtain

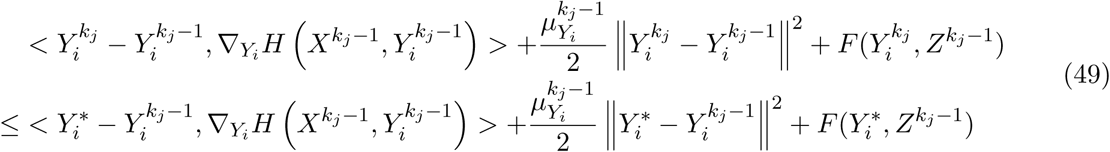

Let *j* → +∞, we get

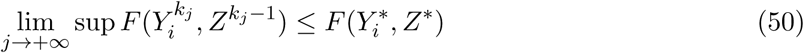

From the fact that *F* is a PLS function, we also have

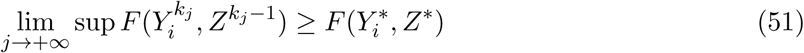

Based on (50) and (51), we know 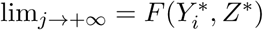. Arguing similarly with *X*, we finally have

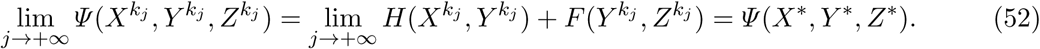

**Condition (iv)**. The function *Ψ* is a semi-algebraic function, which automatically satisfies the Kurdyka-Lojasiewicz property [].

asdfasd

□

### C GPALM Algorithm for NetREX-CF

In this section, we provide the detailed derivation for updating *A, S, x*_*i*_, and *y*_*i*_ used in the GPALM algorithm for NetREX-CF.

#### C.1 Update A

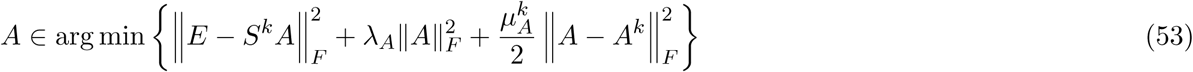

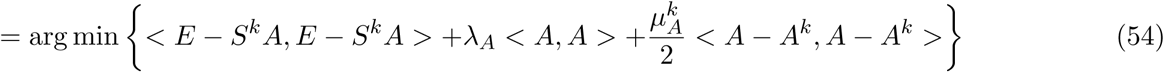

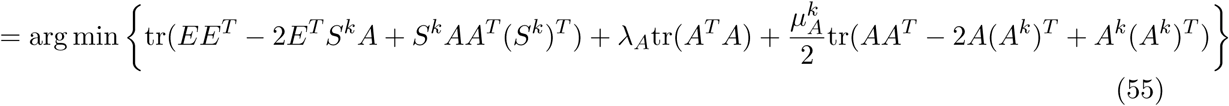

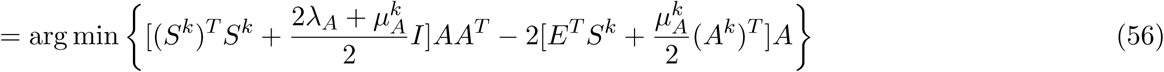

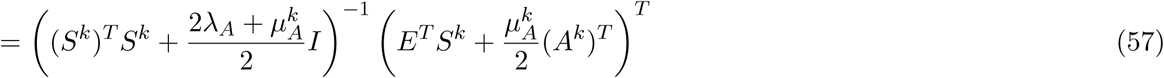

If we complete the square and disregard the constant we get

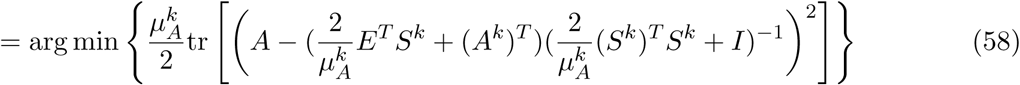

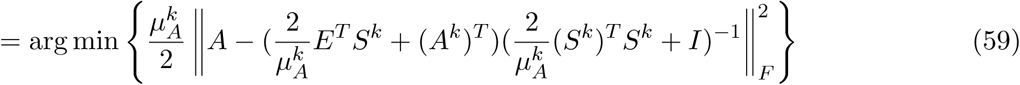

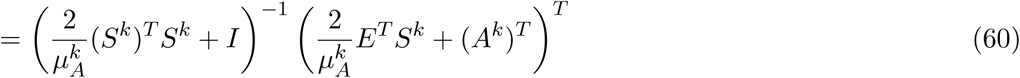

The derivative ∇_*A*_*H*(*A*^*k*^, *S*^*k*^) can be computed by 2((*S*^*k*^)^*T*^ *S*^*k*^*A*^*k*^ + *λ*_*A*_*A*^*k*^ − (*S*^*k*^)^*T*^ *E*), which is Lipschitz continuous with Lipschitz constant 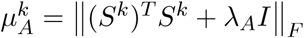.

#### C.2 Update S

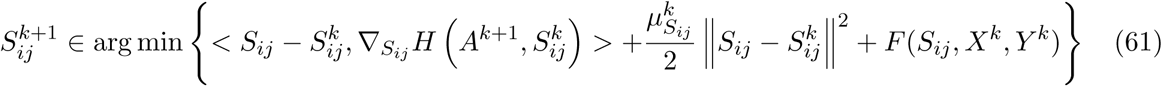

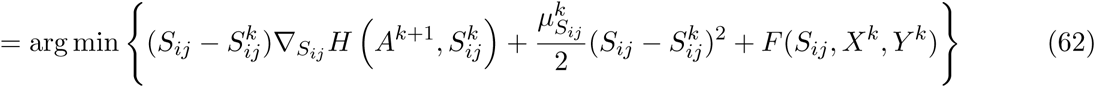

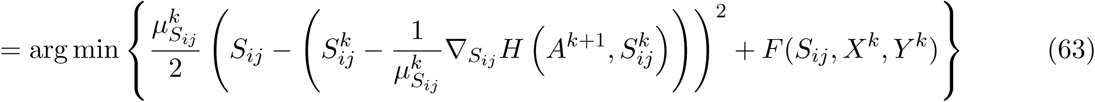

The last equation comes from completing the square and disregarding the constant terms. Now set 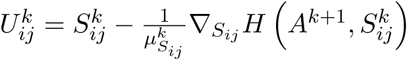 and we have:

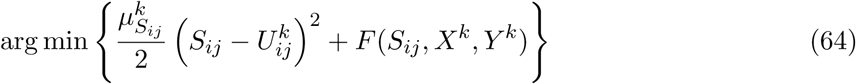

Put *F*(*S, X*^*k*^, *Y* ^*k*^) in and after some algebra.

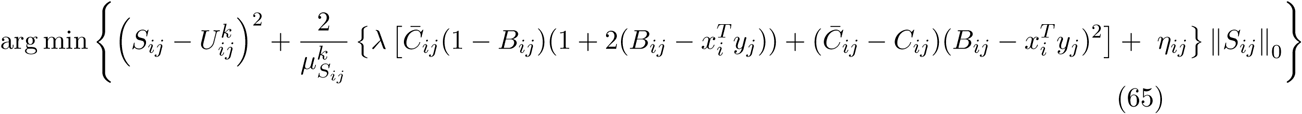

By setting 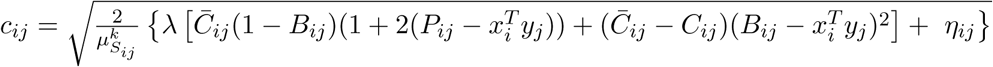, we obtain the hard thresholding problem:

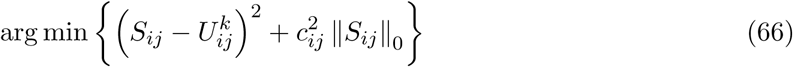

which has solution

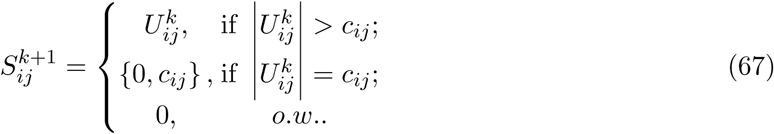

The derivative ∇_*S*_*H*(*A*^*k*+1^, *S*^*k*^) can be computed by 2((*S*^*k*^ *A*^*k*+1^ (*A*^*k*+1^)^*T*^ + *λ*_*S*_*S*^*k*^ − *E*(*A*^*k*+1^)^*T*^), which is Lipschitz continuous with Lipschitz constant 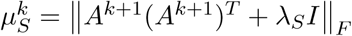.

#### C.3 Update X

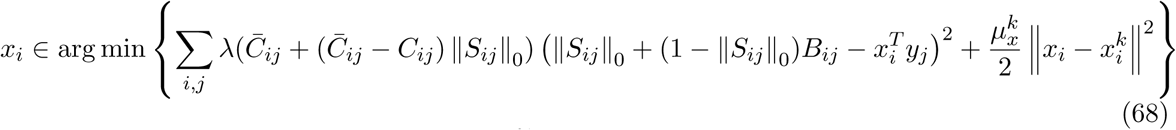

Let 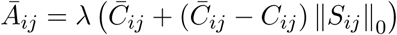 and *Ã* be the diagonal matrix with the values *Ā*_*i*1_, *Ā*_*i*2_, ..*Ā*_*im*_ on the diagonal. Let 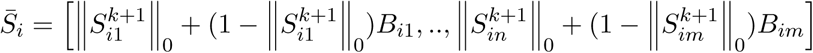. We will show that

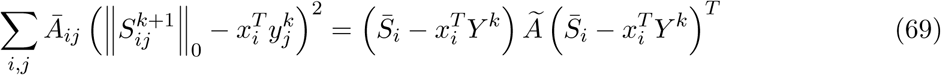

We have

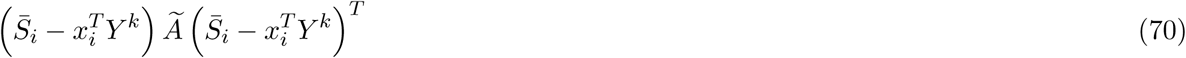

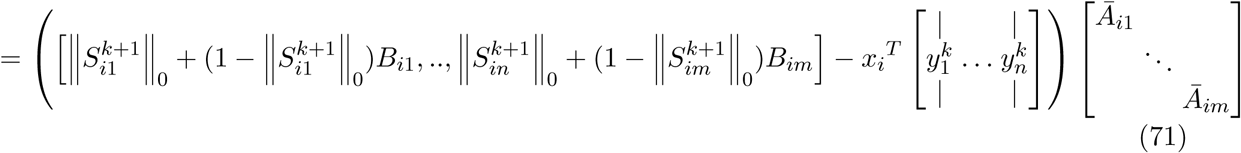

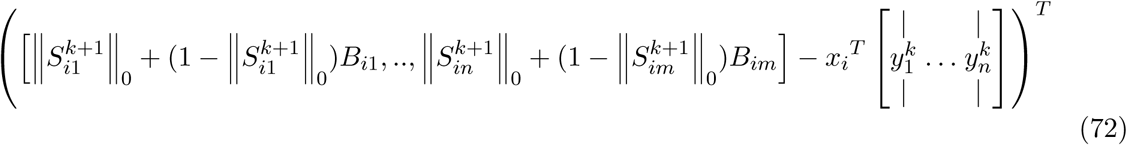

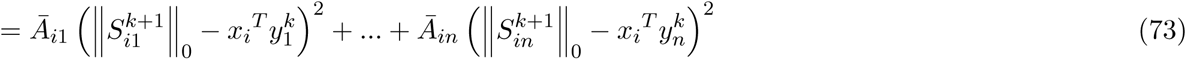

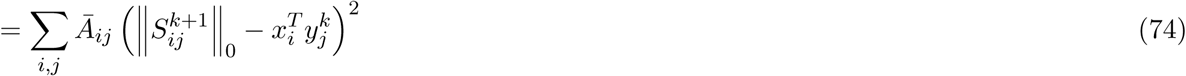

We can now rewrite the problem in terms of the matrix formulation:

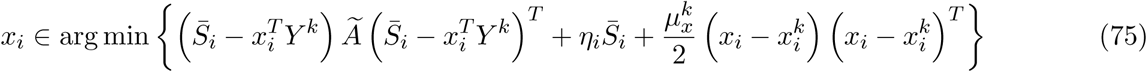

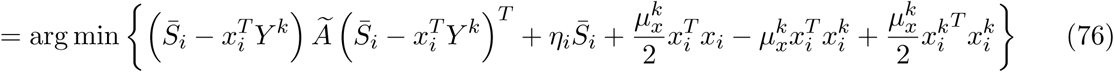

From here we simplify the problem by expanding the first term and removing the constant terms.

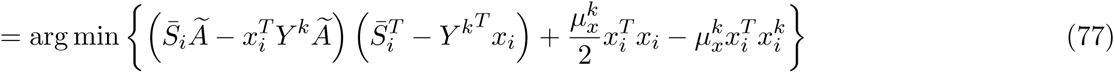

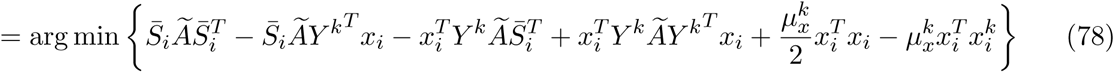

The first term is a constant so we ignore it. Because the third term is a number it is equivalent to its transpose, so the second and third term are the same. We also note that 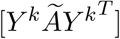 has dimension *h* × *h*, and the second to last term may be rewritten 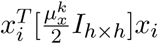.

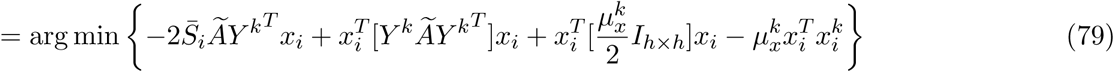

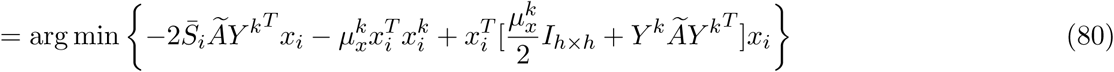

We now use the fact that 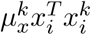 is a number and it is equivalent to its transpose,

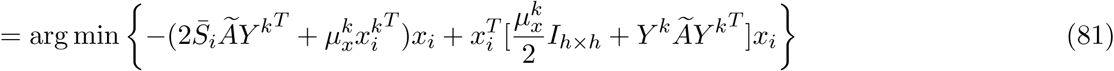

Finally, we define 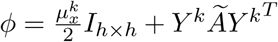 and 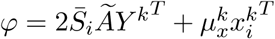, and the problem becomes a Quadratically Constrained Quadratic Program (QCQP) of the form

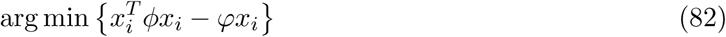

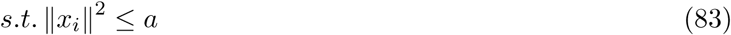

Since we know this problem can be solved, we leave the rest to the CVXPY python package. Lastly, for 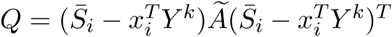 we find the partial gradient

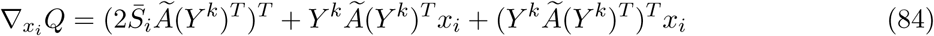

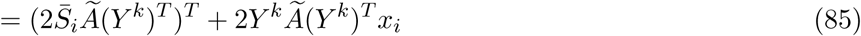

which is Lipschitz continuous with Lipschitz constant 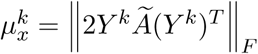.

#### C.4 Solution for Y

Now, for Y, let *Ā*_*ij*_ and *Ã* be the same, but *Ã* will have the values *Ā*_1*j*_, *Ā*_2*j*_, ..*Ā*_*nj*_ on the diagonal. Let *X*^*k*+1^ be a *h* × *n* dimensional matrix where the columns are composed of the vectors 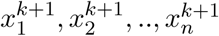 and 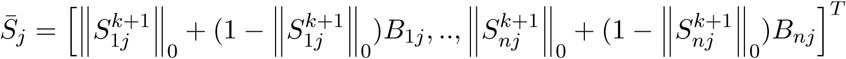. We show that the matrix formulation for this problem by proving the following:

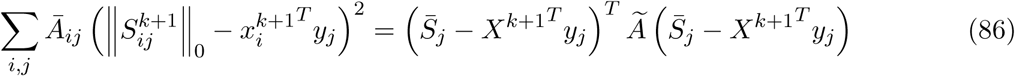

We have

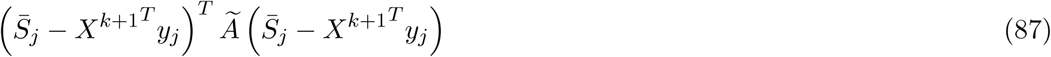

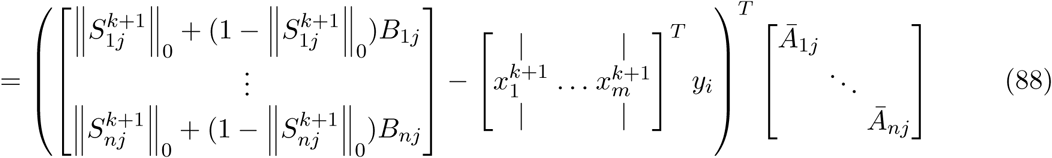

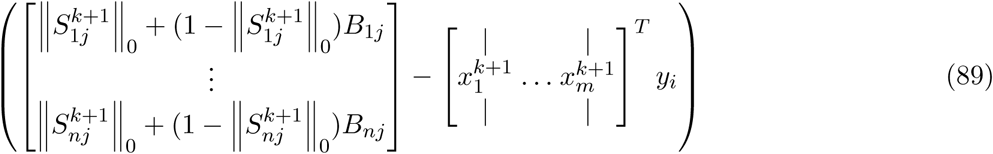

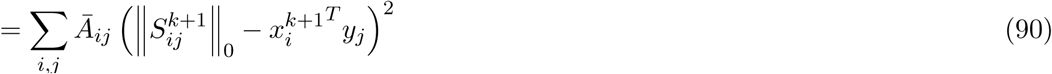

We follow the same procedure to solve this problem

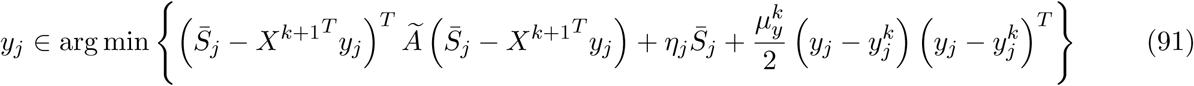

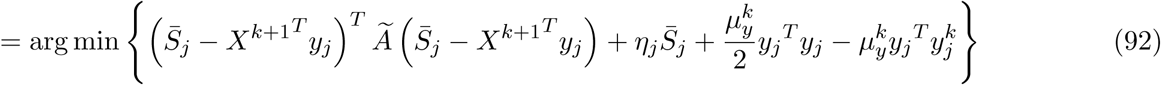

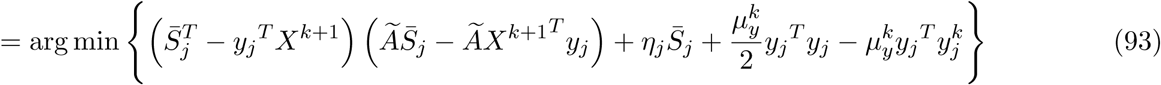

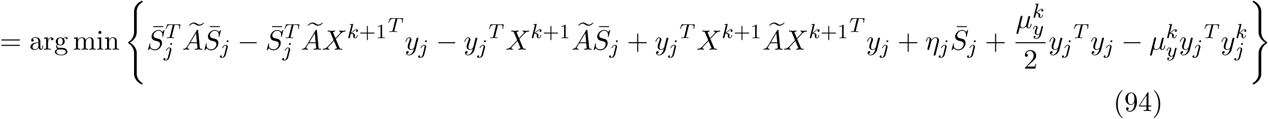

Disregard constants

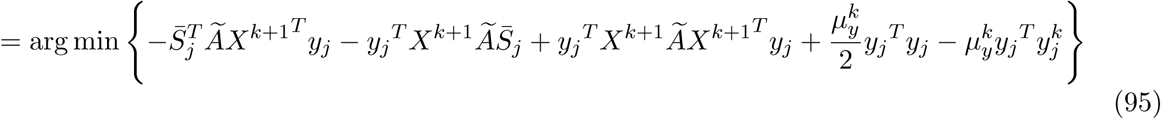

Once again, we can take the transpose of the second term and last term,

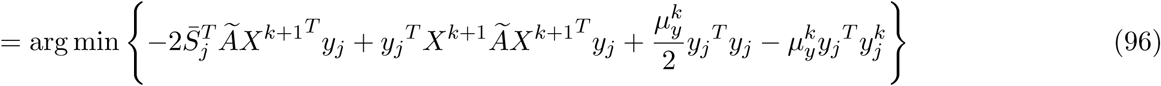

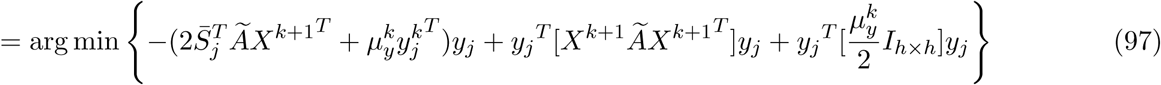

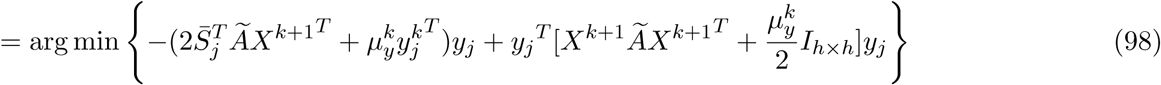

Letting 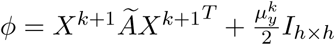 and 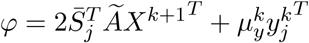 gives us the QCQP

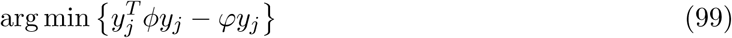

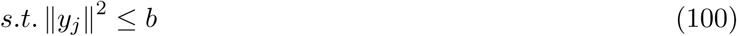

which we solve in CVXPY using the same function. To finish the problem we take 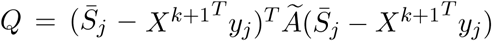 and find the partial gradient

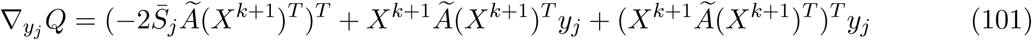

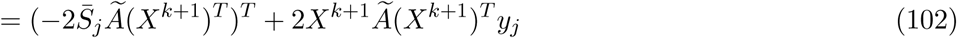

which is Lipschitz continuous with Lipschitz constant 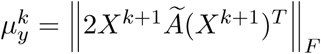.

### D Parameter Selection

In this section, we introduce how we select parameters for the competing algorithms.

#### D.1 Paramter Selection for PriroSum

PriorSum constructs a predicted GRN by summing over weights from all prior networks *P* = {*P*^1^, *…, P*^*d*^}. Therefore, PriorSum builds a GRN 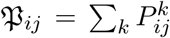 and does not need to select any parameters.

#### D.2 Parameter Selection for LassoStARS

LassoStARS [4] is the latest version of Inferelator, it takes an unweighted prior and gene expression data as input. Because LassoStARS needs an unweighted prior network and the prior networks we have are weighted prior networks, we choose different cutoffs to construct prior networks for LassoStARS. We generate prior networks by assigning each gene the top *N* TFs based on the 𝔓_*ij*_. For *N*, we set *N* = {10, 20, 30, 40} and we find that *N* = 10 performs the best and report the results in Fig. 2. For other parameters used in LassoStARS, LassoStARS proposed a way to select the optimal parameters, therefore, we do not need to select other parameters.

#### D.3 Parameter Selection for MerlinP

For reconstructing the GRN for yeast, MerlinP [5] use the same prior networks and gene expression to build a GRN and reported in the repository https://github.com/Roy-lab/merlin-p. We directly download the GRN they build and compared it with other methods.

#### D.4 Parameter Selection for NetREX

NetREX [2] is similar to LassoStARS, taking an unweighted prior and gene expression as input. So similarly, we generate prior networks for NetREX by assigning each gene the top *N* TFs based on the P_*ij*_. We set *N* = {10, 20, 30, 40} and we find that *N* = 20 performs the best and report the results in Fig. 2. For the other parameters, we selected based on the suggestion provided in https://github.com/ncbi/NetREX.

#### D.5 Parameter Selection for CF

We input CF [16] with 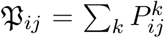. The dimension of the hidden feature vector we set it to be 100, 200, and 300. The regulation term used by CF is set to be 0.1, 1, 10, 100. We try all those combination and report the result with the best performance.

#### D.6 Parameter Selection for NetREX-CF

Based on the formulaiton of NetREX-CF (8), we know that we need to select *h, λ*_*A*_, *λ*_*S*_, *η*_*ij*_, *λ*, and 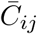. *h* is the dimension of the hidden feature vector. We find that *h* = {100, 200, 300} does not change the performance much. For computational consideration, we set *h* = 100. Because *λ*_*A*_ and *λ*_*S*_ are used as standard regulation to avoid over-fitting, we set *λ*_*A*_ = 1.0 and *λ*_*S*_ = 1.0 by default. We introduce the selection of *η*_*ij*_ and 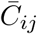 in the following subsection.

**Selection of *η***_***ij***_ We need to make sure *F*(*S, X, Y*) is lower semi-continuous. We can first simplify the equation into

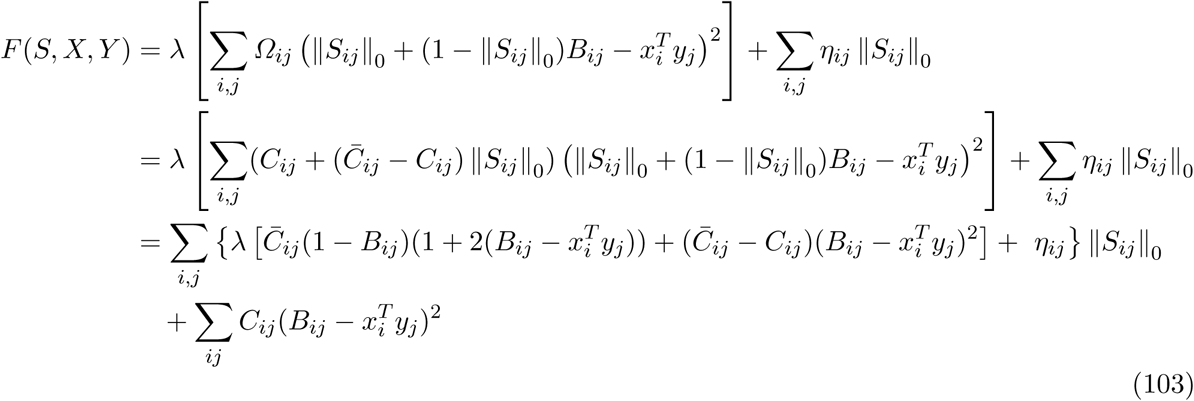

*F*(*S, X, Y*) is lower semi-continuous when the parameter before ‖*S*_*ij*_‖_0_ in the above equation is larger than 0. After several manipulation, we find out we need to set *η*_*ij*_ as following to make *F*(*S, X, Y* lower semi-continuous.

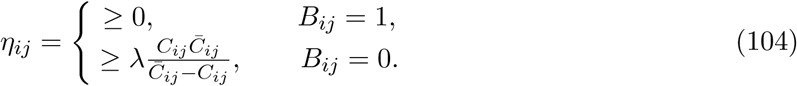

**Selection of** 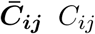 is the penalty when we want to use 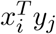 to learn *B*_*ij*_ = 1. Similarly, 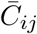 is the penalty when we want to use 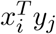 to learn ‖*S*_*ij*_‖_0_ = 1. There are two siutations. First, when ‖*S*_*ij*_‖_0_ = 1 and *B*_*ij*_ = 1, meaning the sparse NCA-based method confirms the edge in the prior, then intuitively, we need to set 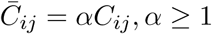. Another situation is that ‖*S*_*ij*_‖_0_ = 1 and *B*_*ij*_ = 0, meaning the sparse NCA-based model confirms an edges recommended by the CF model but not appeared in the prior networks. For this case, we set 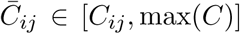, where max(*C*) is the largest element in penalty matrix *C*. In sum, 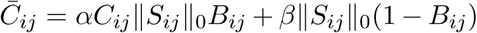, where *α* ≥ 1 and *β* ∈ [*C*_*ij*_, max(*C*)].

**Consensus of Different Parameter Selections** As explained in the previous, for *η*_*ij*_ and 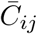, we know the range of these parameters but do not know the exact optimal values. For reconstructing GRN for the yeast experiment, we set

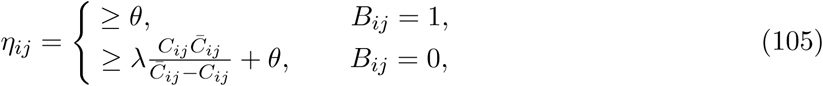

where *θ* = {0.1, 0.5, 1, 2}. And 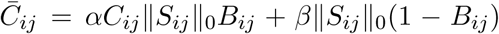, where *α* = {1, 2, 3, 10} and *β* = 10, 20, 30, 40. For different set of parameters, we get a GRN and we get a set of GRNs G = {*G*^1^, *…*}, where *G*^*i*^ = *X*^*T*^*Y* after applying all theses parameters. The final perdition is the average overall predictions 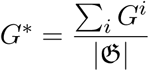.

